# Examining Effects of Schizophrenia on EEG with Explainable Deep Learning Models

**DOI:** 10.1101/2022.05.26.493659

**Authors:** Charles A. Ellis, Abhinav Sattiraju, Robyn Miller, Vince Calhoun

## Abstract

Schizophrenia (SZ) is a mental disorder that affects millions of people globally. At this time, diagnosis of SZ is based upon symptoms, which can vary from patient to patient and create difficulty with diagnosis. To address this issue, researchers have begun to look for neurological biomarkers of SZ and develop methods for automated diagnosis. In recent years, several studies have applied deep learning to raw EEG for automated SZ diagnosis. However, the use of raw time-series data makes explainability more difficult than it is for traditional machine learning algorithms trained on manually engineered features. As such, none of these studies have sought to explain their models, which is problematic within a healthcare context where explainability is a critical component. In this study, we apply perturbation-based explainability approaches to gain insight into the spectral and spatial features learned by two distinct deep learning models trained on raw EEG for SZ diagnosis for the first time. We develop convolutional neural network (CNN) and CNN long short-term memory network (CNN-LSTM) architectures. Results show that both models prioritize the T8 and C3 electrodes and the δ- and γ-bands, which agrees with previous literature and supports the overall utility of our models. This study represents a step forward in the implementation of deep learning models for clinical SZ diagnosis, and it is our hope that it will inspire the more widespread application of explainability methods for insight into deep learning models trained for SZ diagnosis in the future.

## I. Introduction

Schizophrenia (SZ) is a mental disorder that affects more than 20 million individuals worldwide [1]. Because of variation in symptoms and a lack of clinically-utilized biomarkers, diagnosis can be challenging, and many studies in recent years have focused on developing automated methods of SZ diagnosis with electroencephalography (EEG) data. Early attempts at automated SZ diagnosis frequently required complicated feature engineering [2], [3], and more recent studies have sought to use more advanced deep learning frameworks to automate feature extraction from raw EEG [1][4]. While these studies do make feature extraction easier, they are limited in their utility for the healthcare domain in that they do not provide insight into how they work and what features they have learned. In this study, we develop two distinct deep learning architectures inspired by existing studies and for the first time apply explainability methods for insight into the key spectral and spatial features associated with SZ diagnosis in raw EEG data.

The diagnosis of SZ is typically based upon symptoms. Symptoms of SZ can include a decline in cognitive function, lack of pleasure, flattening of emotion, delusions, hallucinations, difficulty concentrating, and disorganized speech [1]. While these symptoms are well-known, they vary from person to person, which can make diagnosis challenging. As such, studies have sought to identify biomarkers of SZ and develop automated SZ diagnostic methods for clinical decision support. Many of these studies have utilized machine learning with EEG data.

Early attempts at automated diagnosis of SZ involved manual feature engineering with traditional machine learning models [2]. However, manual feature engineering can be challenging, and later studies have begun to apply deep learning frameworks to raw EEG to enable automated feature extraction and evade the difficulty of manual feature engineering [1][4]. While this represents a step forward in ease of feature extraction, it also makes explainability more challenging. Existing deep learning studies using raw EEG have not provided any insight into the features that they extracted and the relative importance of those features. This is problematic given that explainability represents a critical component of any use of deep learning models in a clinical setting [5].

In this study, we apply explainability approaches that provide insight into the importance of spectral and spatial features learned by neural networks trained on raw EEG data for the first time. Using architectures from previous studies as a starting point, we develop two highly distinct architectures: a convolutional neural network (CNN) and a combination of a CNN and long short-term memory network (CNN-LSTM). We then apply our explainability methods, examine which features are highly important to both architectures, and compare our findings with existing literature. Our study represents the first application of explainability methods to neural networks trained on raw EEG for SZ diagnosis and a step towards the clinical implementation of deep learning for SZ diagnosis.

## II. Methods

### A. Data and Preprocessing

We utilized a scalp EEG dataset originally presented in [6]. The dataset contains 47 individuals with SZ and 54 healthy controls (HCs). Data collection was approved by the Hartford hospital IRB, and all participants gave written, informed consent. During recording, 64 electrodes placed in the standard 10-20 format were used, but like previous studies [1][4], we only used 19 electrodes (i.e., Fp1, Fp2, F7, F3, Fz, F4, F8, T3, C3, Cz, C4, T4, T5, P3, Pz, P4, P6, T6, O1, and O2). Recordings were performed during a 5-minute resting state at 1000 Hz. Prior to our analyses, we downsampled the data to 250 Hz, and separated it into 25-second epochs. We then randomly upsampled SZ samples to ensure balanced representation of both SZ and HC data. All samples were then detrended by fitting to a polynomial basis of degree 5 [7]. We removed samples that had amplitudes greater than or less than the 99.99^th^ and 0.01^st^ percentiles, respectively. We then applied z-score normalization for each remaining sample along the channel axis. Our final dataset consisted of 5765 HC and 5764 SZ samples.

### B. Model Development

We developed two distinct deep architectures in this study. We trained both architectures using a 10-fold cross-validation strategy where 80%, 10%, and 10% of the data was the train, validation, and test set, respectively. Data was split by subject rather than by sample, ensuring the model would evaluate complete subject data in the test set that were unseen in the training set. Prior to training, we applied data augmentation on the training data set for each fold. We first generated 3 copies of the training data. We applied Gaussian noise with a mean (µ) of 0 and standard deviation (σ) of 0.7 to one copy. We applied, to the best of our knowledge, a novel data augmentation technique in the form of sinusoidal noise with frequency of 2 Hz and amplitude of 0.3 to another copy. We left the final copy untouched and concatenated all copies to form our augmented training data. We then performed the training process with a CNN and CNN-LSTM.

Figure 1 shows our CNN architecture. It was inspired by an existing architecture with an added emphasis on generalization [1]. To this end, batch normalization and MaxNorm regularization were used. Figure 2 shows our CNN-LSTM architecture. It was inspired by a previous architecture [4]. We used batch normalization in this architecture as well. We also used a weighted loss to account for potential class imbalances caused by the train-validation-test split.

**Fig 1.**
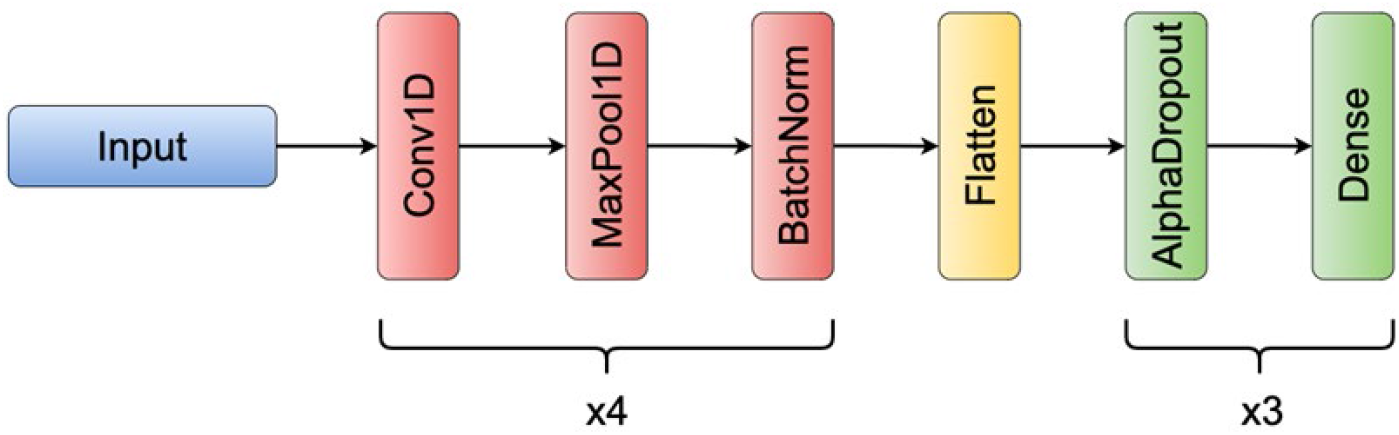
CNN Architecture. There are 4 1D convolution (Conv1D) layers in the convolution block. The first Conv1D layer has 5 filters of length 10 and a stride of 1. The second and third Conv1D layers have 10 filters of length 10 and strides of 1. The fourth Conv1D layer has 15 filters of length 5 and a stride of 1. Each Conv1D layer is followed by Max Pooling layer (pool size of 2, stride of 2) and Batch Normalization layers. The classifier portion of the CNN contains 3 Dense layers with 64 nodes, 32 nodes, and 1 node. All Dense layers are preceded by an AlphaDropout layer with a dropout rate of 0.5. All Conv1D and Dense layers have Exponential Linear Unit (ELU) activations, He Normal weight initialization, and MaxNorm regularization with a maximum value of 1. The final Dense layer has a Sigmoid activation and Glorot Normal weight initialization.

**Fig 2.**
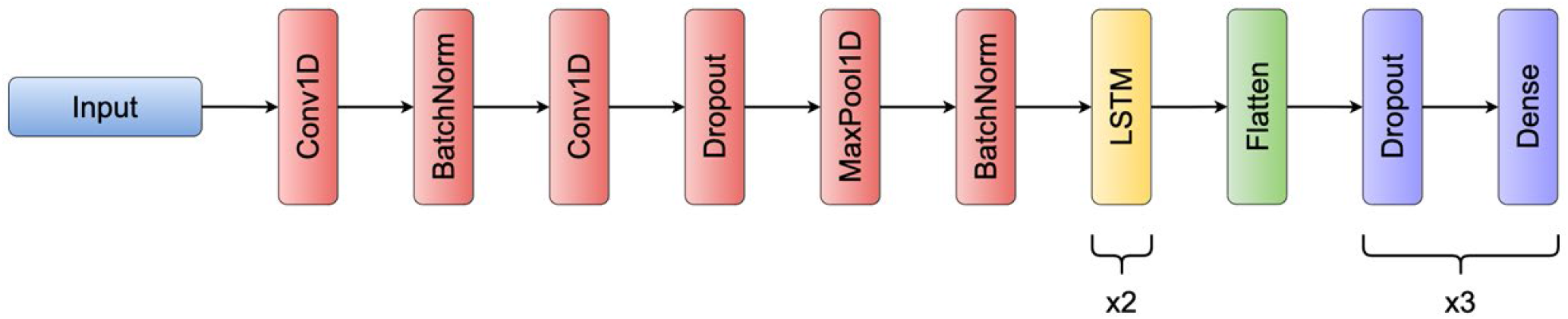
CNN-LSTM Architecture. The CNN-LSTM has 2 convolution layers (both have 64 filters of length 3, stride of 1). All dropout layers have a dropout rate of 0.5. The architecture has 1 max pooling layer (pool size of 2, stride of 1). The first LSTM layer has 128 units, and the second has 64 units. Both LSTM layers return full sequences of time steps. The classifier portion of the CNN-LSTM following the flatten layer contains 3 dense layers with 128 nodes, 64 nodes, and 1 node. All Conv1D and dense layers have a Rectified Linear Unit (ReLU) activation and He Normal weight initialization. The final dense layer has a sigmoid activation and Glorot normal weight initialization. Both LSTM layers have a hyperbolic tangent (tanh) activation with Glorot uniform weight initialization.

For both models, we used binary cross-entropy loss and an Adam optimizer (learning rate = 0.001). We trained the models for 50 epochs with a batch size of 128 and shuffling after each epoch. Across epochs, we tracked accuracy (ACC), sensitivity (SENS), specificity (SPEC), and balanced ACC (BACC) to quantify performance. We used a checkpoint approach to select the model from the epoch with the maximal BACC on the validation set and used that model for testing. predicted on the test set using a checkpointed during training. We calculated the μ and σ of the listed metrics across folds.

### C. Explainability Approaches

We employed two explainability approaches in our study. In our first approach, we ablated individual EEG channels, and in our second approach, we ablated specific frequency bands.

#### 1) Spatial Explainability

For insight into the importance of the spatial features learned by the models, we used a channel-based perturbation approach with the following steps: (1) We calculated the BACC of the model on the test data. (2) We iterated through each of 19 channels and ablated each channel individually across all samples with sinusoidal noise with frequency of 60 Hz and amplitude of 0.1. (3) We measured the percent change in BACC after each perturbation. Our approach mimicked the line-related noise commonly found in EEG and that has been used in multimodal explainability [8].

#### 2) Spectral Explainability

For insight into the importance of the spectral features learned by the models, we employed a perturbation approach involving multiple steps: (1) We calculated the BACC of the model on the test data. (2) We converted each sample to the frequency domain with a Fast Fourier Transform (FFT). (3) We randomly permuted spectral coefficients within each frequency band across samples and channels such that all coefficients belonging to a particular frequency band for a sample and subject were substituted for the coefficients in a separate sample and subject. (4) We converted the perturbed samples back to the time domain with an inverse FFT, and (5) we calculated the percent change in BACC after perturbation. At each iteration we perturbed a different frequency band in Hertz (Hz): *δ* (0 – 4 Hz), *θ* (4-8 Hz), α (8 – 12 Hz), β (12 – 25 Hz), γ_lower_ (25 – 55 Hz), and γ_upper_ (65 – 125 Hz).

## III. Results and Discussion

In this section, we describe and discuss our model performance and explainability results.

### A. Model Performance

Table 1 shows the μ and σ of the ACC, SENS, SPEC, and BACC of our classifiers. Both classifiers performed well above chance-level, obtaining performance metrics at or above 70%. Overall, the performance of our CNN was higher than that of the CNN-LSTM. Additionally, the mean SPEC of the CNN was somewhat higher than its mean SENS, while the mean SENS of the CNN-LSTM was slightly higher than its mean SPEC. The mean SENS of our CNN was slightly higher than the CNN in [1], and the mean SPEC and ACC of our CNN were slightly lower. The performance of our models was far below the performance of the CNN-LSTM models in [4]. However, that study used a cross-validation approach that allowed samples from the same subjects to be assigned to training, validation, and test sets simultaneously. Given the dependencies that exist within time-series, that approach can artificially inflate performance, and test performance no longer actually indicates the generalizability of the patterns learned by the model. Our cross-validation approach enabled a better understanding of the generalizability of the models.

**TABLE I.**
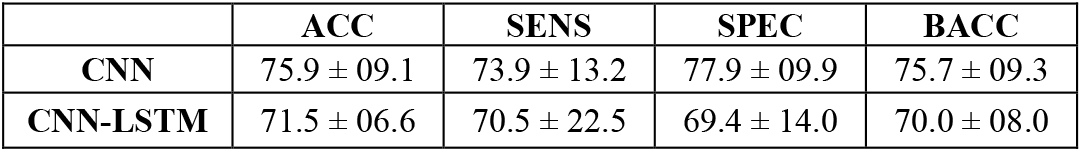
Classification Performance Results

### B. Spatial Explainability

Figure 3 shows the effects of ablating individual channels on the performance of our classifiers. Interestingly, two of the top five most important channels for each architecture were shared (i.e., T8 and C3), and the importance of these channels fits with existing literature. T8 is located near the temporal lobe, which has been found to have increases in δ activity in SZs [9], and C3 has previously been found to have less complexity in SZs than HCs [10]. Nevertheless, the two classifiers still seemed to have some strong differences in the channels that they prioritized that are possibly attributable to how the two architectures treat temporal dependencies in extracted features differently.

**Fig. 3.**
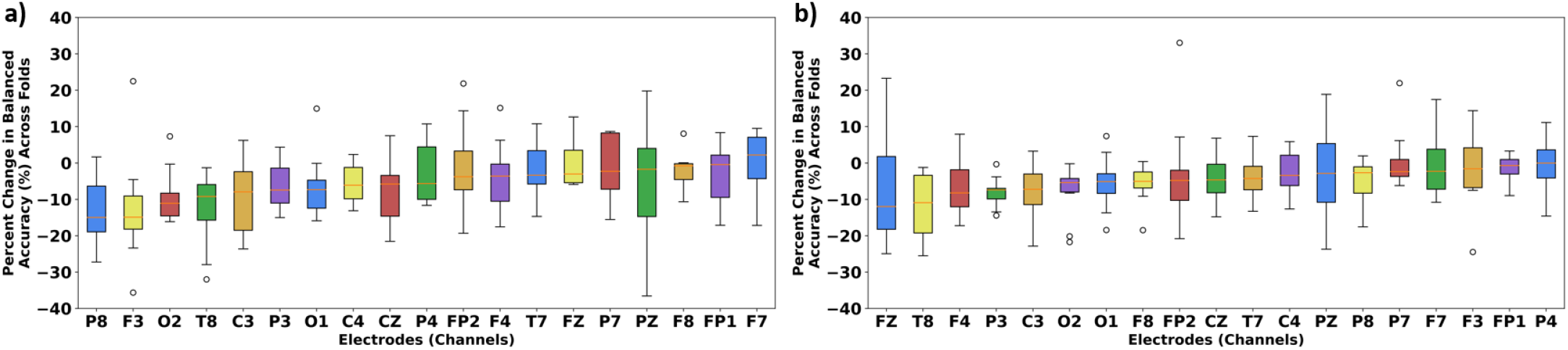
Plots of Spatial Importance. Panels a) and b) show the importance of each channel to the CNN and CNN-LSTM, respectively. Channels are arranged on the x-axis in order of importance. The percent change in BACC following perturbation across folds is shown on the y-axis.

### C. Spectral Explainability

Figure 4 shows the effects of perturbing different frequency bands on the performance of our classifiers. For both classifiers, θ, α, and β were not considered to be very important, though θ did have some impact upon the CNN. Instead, δ, γ_lower_, and γ_upper_ were important. The CNN most prioritized γ_upper_ followed by δ and γ_lower_. In contrast, the CNN-LSTM most prioritized δ followed by γ_upper_ and γ_lower_. Our results are similar to those found in previous studies that have identified key differences between HCs and SZs in the δ- and γ-bands [11].

**Fig. 4.**
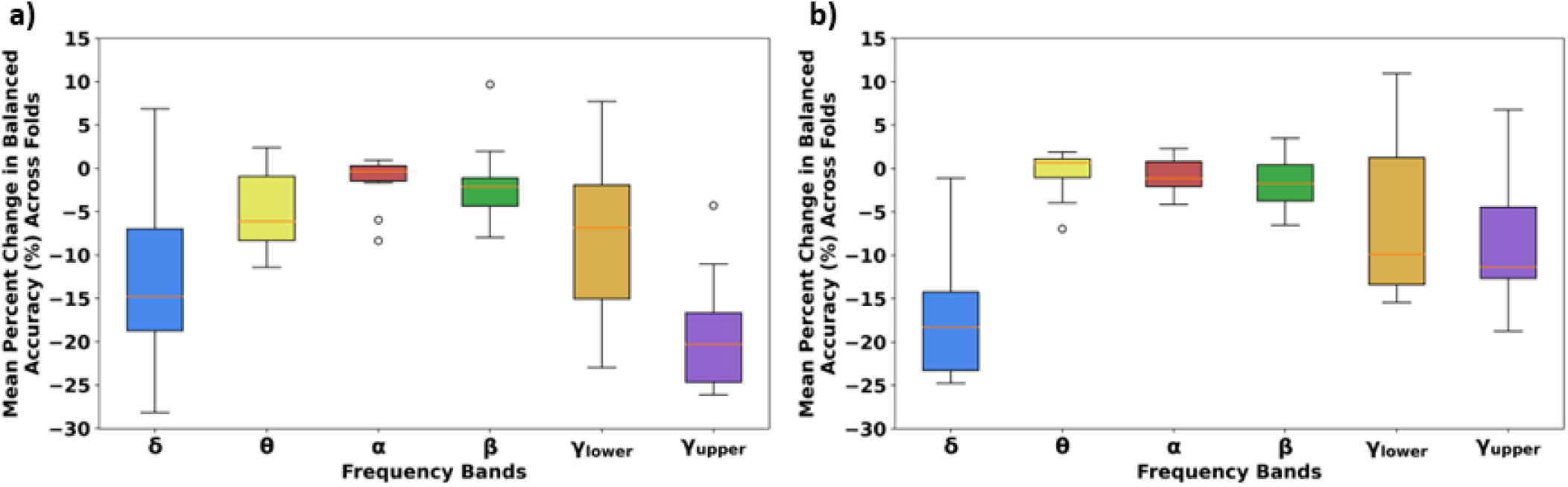
Plots of Spectral Importance. Panels a) and b) show the importance of the canonical frequency bands for the CNN and CNN-LSTM, respectively. The frequency bands are arranged on the x-axis, and the percent change in BACC following perturbation across folds is shown on the y-axis. In this figure, δ, θ, α, β, γ_lower_, and γ_upper_ are shown in blue, yellow, red, green, orange, and purple, respectively.

### D. Limitations and Next Steps

In this study, we examined the importance of each channel and each frequency band across all channels. In future studies, it would be interesting to examine the effect of perturbing each frequency band within each channel. It is also possible that gradient-based methods could be used to gain insight into the spatial features learned by the CNN [12] or that an attention mechanism could be integrated into the CNN-LSTM. In these cases, it would be possible to gain insight into the temporal effects of SZ on EEG activity by examining the temporal distribution of importance. It would also be interesting to develop a more interpretable classifier to gain insight into the waveforms that might be useful for SZ diagnosis [13], [14].

## IV. Conclusion

Our study represents the first application of explainability methods to the domain of automated SZ diagnosis with raw EEG. We adapted two highly distinct deep learning architectures to perform SZ diagnosis and then applied two explainability approaches for insight into the spatial and spectral features learned by the models. Both models prioritized the T8 and C3 electrodes, and we found that the δ- and γ-bands were important to the performance of the classifiers. The findings of our explainable deep learning architectures are corroborated by existing literature, and our study represents a step towards the implementation of automated SZ diagnosis in a clinical setting.

## Acknowledgment

We thank those who collected the EEG data used in this study.

